# Age-related neural dedifferentiation for individual stimuli: An across-participant pattern similarity analysis

**DOI:** 10.1101/2021.08.27.457968

**Authors:** Joshua D. Koen

## Abstract

Age-related neural dedifferentiation – reductions in the regional specificity and precision of neural representations – is proposed to compromise the ability of older adults to form sufficiently distinct neural representations to support episodic memory encoding. The computational model that spurred investigations of age-related neural dedifferentiation initially characterized this phenomenon as a reduction in the specificity of neural patterns for individual items or stimuli. Most investigations have focused on reductions in neural differentiation for patterns of neural activity associated with category level information, such as reduced neural selectivity between categories of visual stimuli (e.g., scenes, objects, and faces). Here, I report a novel across-participant pattern similarity analysis method to measure neural distinctiveness for individual stimuli that were presented to participants on a single occasion. Measures of item level pattern similarity during encoding showed a graded positive subsequent memory effect in younger, with no significant subsequent memory effect in older adults. These results suggest that age-related reductions in the distinctiveness of neural patterns for individual stimuli during age differences in memory encoding. Moreover, a measure of category level similarity demonstrated a significant subsequent memory effect associated with item recognition (regardless of an object source memory detail), whereas the effect in older was associated with source memory. These results converge with predictions of computational models of dedifferentiation showing age-related reductions in the distinctiveness of neural patterns across multiple levels of representation.

## Introduction

Aging is associated with declines in numerous cognitive domains, most notably in the domain of episodic memory—memory for unique events from our past (Drag & Bieliauskas, 2010; Salthouse, 2019). The present study focuses on the contribution of neural dedifferentiation to age differences in encoding processes that support episodic memory. Neural dedifferentiation refers to the finding that neural representations become less distinct with increasing age, which in turn compromises memory and cognition in older adults (Koen et al., 2020; Koen & Rugg, 2019; Li et al., 2001; Li & Rieckmann, 2014). Neural dedifferentiation might increase the likelihood that memory traces that are formed during encoding are not sufficiently distinct to support retrieval of specific details (Craik & Rose, 2012), which might explain age-related deficits in recollection (for reviews, see Koen & Yonelinas, 2014; Schoemaker et al., 2014). Critically, evidence from computational modelling (Li et al., 2001; Li & Rieckmann, 2014) and the application of multivariate analysis techniques to human neuroimaging data (Kobelt et al., 2021; for reviews, see Goh, 2011; Koen et al., 2020; Koen & Rugg, 2019) indicate that neural dedifferentiation occurs over multiple levels of neural representation to include neural representations for categorical information and item-specific information. Understanding how age-related neural dedifferentiation for these different levels of representation contribute to memory will shed light on the factors contributing to age-related declines in memory. The present study examines this issue using across-participant multivariate pattern similarity analysis of fMRI data.

Research on age-related neural dedifferentiation has mainly focused on age differences in category level neural representations (Abdulrahman et al., 2017; Carp et al., 2011; Chamberlain et al., 2021; Kleemeyer et al., 2017; Koen et al., 2019; D. C. Park et al., 2004; J. Park et al., 2012; Payer et al., 2006; Srokova et al., 2020; T. H. Wang et al., 2016; Zebrowitz et al., 2016; for reviews, see Koen & Rugg, 2019; Koen et al., 2020). Category level representations can be quantified using univariate measures, such as the neural differentiation index (Koen et al., 2019; Srokova et al., 2020; Voss et al., 2008; Zebrowitz et al., 2016), and with multivariate analytic techniques, such as multivariate classification and pattern similarity analysis (Chamberlain et al., 2021; Kleemeyer et al., 2017; Kobelt et al., 2021; J. Park et al., 2010; Srokova et al., 2020; Trelle et al., 2019). Of relevance here, pattern similarity analysis quantifies category level neural representations by contrasting the similarity (i.e., correlation) of the spatial profile of neural activation between trials belonging to the same stimulus category (e.g., scenes; within-category similarity) with trials that belong to a different stimulus category (e.g., faces or objects; between-category similarity). I will refer to this measure as *category level similarity*. Research using pattern similarity analyses (e.g., Chamberlain et al., 2021; Kleemeyer et al., 2017; J. Park et al., 2010; Srokova et al., 2020), and other approaches (e.g., Koen et al., 2019; D. C. Park et al., 2004; Payer et al., 2006; Voss et al., 2008; Zebrowitz et al., 2016), has provided robust evidence for age-related neural dedifferentiation of category level representations. Category level dedifferentiation is also associated with memory performance whereby individuals with less distinctive category level representations, regardless of age, perform more poorly on recognition memory tests (Kobelt et al., 2021; Koen et al., 2019; Srokova et al., 2020). Overall, the existing data provides strong evidence that category level representations are less distinct in older adults, and that such representations also play a role in episodic memory regardless of age.

There have been comparatively fewer studies investigating whether item-specific neural representations differ between young and older adults and if they relate to memory. The most common approach used to measure item-specific representations is to compare pattern similarity across repetitions of the same stimulus (Kobelt et al., 2021; St-Laurent et al., 2014; Trelle et al., 2019; Xue et al., 2010; Zheng et al., 2018; for a related multivariate approach, see Bowman et al., 2019). Measures of *item level similarity* are obtained for each unique stimulus by contrasting the similarity across repetitions of the exact same item (within-item similarity) with similarity across different items belonging to the same category (within-category similarity). There have been less consistent findings with respect to age differences in measures of item level similarity and their relationship to episodic memory (for discussion, see Koen & Rugg, 2019). St-Laurent and colleagues (2014) observed null age differences for measures of item level similarity during encoding. In contrast, results from other studies have shown that measures of item level similarity were reduced in older adults relative to young adults (Kobelt et al., 2021; Trelle et al., 2019; Zheng et al., 2018). Moreover, only two of these studies (Kobelt et al., 2021; Zheng et al., 2018) directly related measures of item level similarity to subsequent memory outcomes. For instance, Kobelt and colleagues reported an age-invariant negative relationship between item level similarity and memory performance, which is like the brain-behavior relationships observed with dedifferentiation of category level neural representations discussed above. Given these mixed findings and relatively limited research on this topic, additional research investigating the relationship between dedifferentiation of item level representations, age, and memory is needed.

The current report investigates age differences in both category level and item level neural representations and how they relate to successful memory encoding. To do so, I conducted a reanalysis of data initially reported by Koen and colleagues (2019). In this study, cognitively healthy young and older adults encoded images of objects and scenes for a subsequent recognition test that measured both item recognition (i.e., old and new images) and source memory (i.e., memory for a semantic judgment performed during encoding). Here, I used a novel across-participant pattern similarity analysis to measure category level and item level similarity and relate these measures to subsequent memory outcomes. There is growing interest in using across-participant analyses, with a particular focus on using the method to measure item-specific patterns of neural activity (Chen et al., 2017; Koch et al., 2020; Xiao et al., 2020; Zadbood et al., 2017; also see Hasson et al., 2004; for a methodological overview, see Q. Wang et al., 2020). This approach can also, in principle, be used to examine patterns of neural activity for category level information (for an across-participant multivoxel classification analysis, see Thakral et al., 2019).

Adopting an across-participant analysis approach to measure item-specific neural activity is potentially advantageous compared to the more common within-participant pattern similarity approaches described above. Specifically, across-participant pattern similarity analyses can be applied to paradigms in which a given stimulus is presented on only one occasion, which can obviate issues associated with the influence of stimulus repetitions on pattern similarity measures (cf. St-Laurent & Buchsbaum, 2019). The across-participant pattern similarity analysis developed here leverages linear-mixed effects models to obtain predicted item-specific and category level neural patterns (see Figure 1). These predicted maps are, in turn, related to observed neural patterns to measure item level and category level similarity. These two measures are then subjected to a subsequent memory analysis to examine their relationship to successful memory encoding.

**Figure 1.**
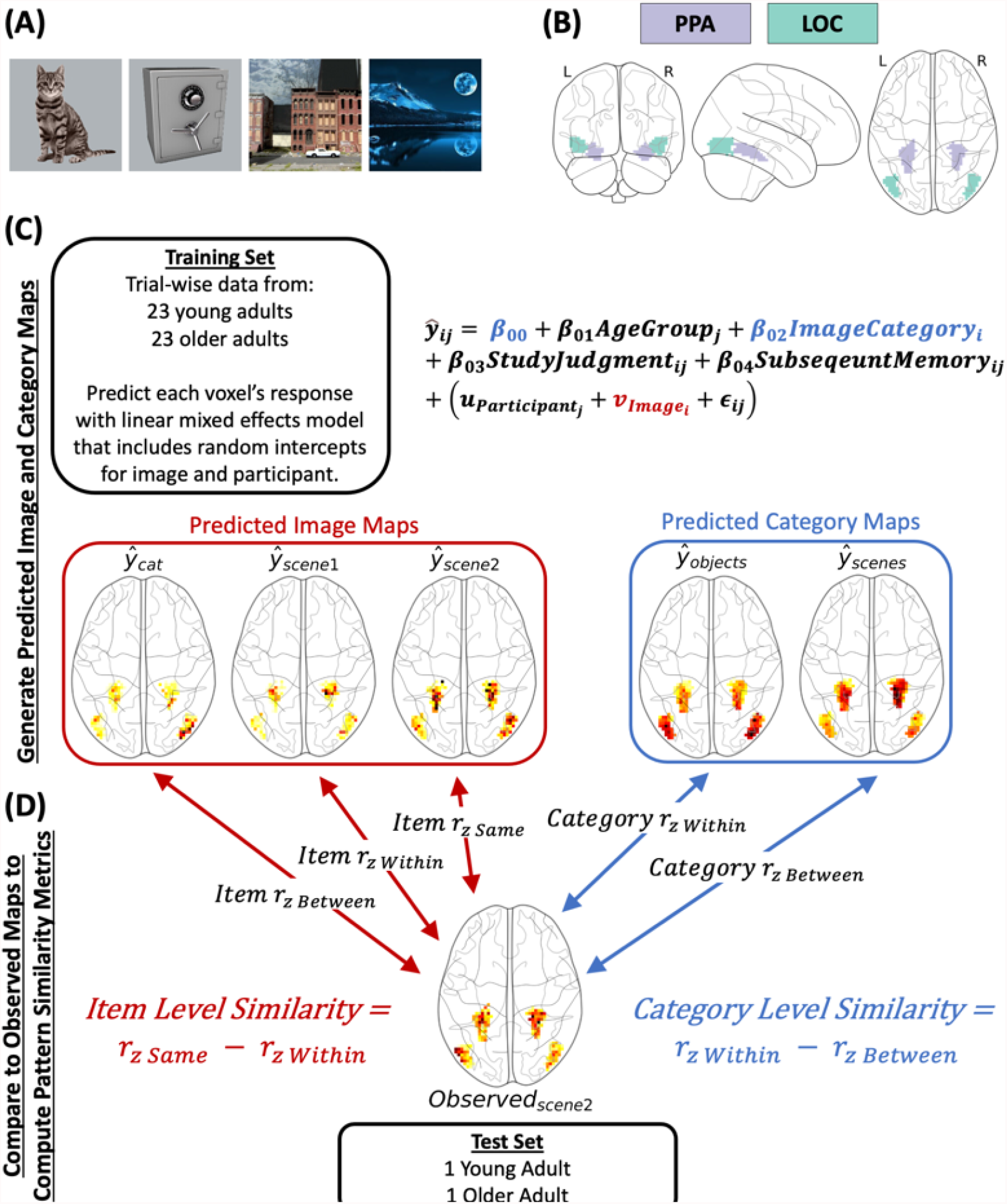
(A) Example stimuli from the experiment. (B) The parahippocampal (PPA) and lateral occipital complex (LOC) regions of interest. The PPA and LOC were combined into a single region of interest for the primary analysis. (C) Visual depiction of the across-participant procedure used to compute predicted image and category maps. The approach was iterative whereby, on each iteration, data from the 23 young and 23 older adults (with yoked stimulus sets; the ‘ training set’) were submitted to a linear mixed effects model with random intercepts for image 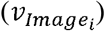 and participant 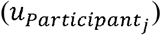 and, of most relevance here, the fixed effect term for image category (*β*_02_). This model was used to predict the response (i.e., beta value) in each voxel in two regions of interest shown in Panel B. The predicted image map for each image (320 total) was created by taking the random intercept coefficient 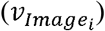 for the image at each voxel (omitting all other terms in the model). The predicted category maps (one for scenes and one for objects) were computed in a similar manner using the model intercept (*β*_00_) and the coefficient for the image category term (*β*_02_). (D) Item and category level similarity were obtained by examining the similarity of the predicted image and category maps to the observed beta maps from the held out young and older adult (i.e., the ‘ test set’). The similarity metric (i.e., the Fisher-*z* transformed Pearson correlation) was used to obtain measures of item level and category level similarity. Item level similarity was defined as the difference in similarity between the observed and predicted image maps for the same image (e.g., scene 2) and the average similarity between the observed image and the predicted images in the same category (e.g., scene 2 with all other scenes). Category level similarity was measured as the difference in similarity between the observed and predicted category map from the same (within) and other (between) category. These difference scores for each were used to control for individual differences in baseline similarity, which is critical in studies comparing pattern similarity measures across groups or conditions (Dimsdale-Zucker & Ranganath, 2018). This procedure comprised 24 iterations so that each young and older adult were held out in the ‘ test set’.

## Methods

### Participants

The data contributing to the present analysis comprised a sample of healthy young (N = 24; mean age = 23 years; range = 18-30; 12 females) and older adults (N = 24; mean age = 69 years; range = 64-75; 12 females) initially reported by Koen and colleagues (2019). Participants included individuals recruited from the University of Texas at Dallas and the greater Dallas metropolitan area. Each participants received monetary compensation ($30/hour) for volunteering in the experiment. All participants were right-handed and reported having normal or corrected-to-normal vision, had no contraindications to MRI scanning, and had no history of cardiovascular disease (other than treated hypertension), diabetes, psychiatric disorder, illness, or trauma affecting the central nervous system, substance abuse, nor self-reported current or recent use of psychotropic medication or sleeping aids. All participants were considered cognitively normal as determined by performance battery of neuropsychological tests. Each participant scored 27 or more on the Mini-Mental State Examination (MMSE; Folstein et al., 1975) and no less than 1.5 standard deviations below age-normalized scores on any one memory measure or on two (or more) non-memory measures (see Supplemental Table 1 for a full reporting of the demographic and neuropsychological test data).

The protocol for this experiment was approved by the Institutional Review Board of the University of Texas at Dallas. All participants provided written informed consent prior to participation.

## Materials and Procedure

The detailed materials and procedure for the experimental memory task are reported in Koen et al. (2019) and will be described here in brief. The stimuli comprised 180 images of scenes (half depicting a rural scene; half depicting an urban scene) and 180 images of single objects (half depicting natural objects; half depicting manmade objects; see Figure 1A). These images were used to create a total of 24 yoked stimulus sets. For each stimulus set, a random selection of 120 scenes and 120 objects were assigned to be studied items (shown during both the study and test phase) with the remaining 60 scenes and 60 objects assigned to be new items in the test phase (i.e., not shown during the study phase). Stimuli were presented using Cogent 2000 software (www.vislab.ucl.ac.uk/cogent_2000.php) implemented in Matlab 2011b (www.mathworks.com).

Participants completed the study phase (under intentional encoding instructions) while undergoing fMRI scanning. The study phase was split across 5 consecutive scanning sessions. The sequence and timing for each trial was as follows: get ready signal (green fixation cross for 500 ms), task cue (red “P?” or “M?” for 500 ms), study image (object or scene for 2000 ms), and white fixation (1750 ms). For each image, participants were instructed to make a pleasantness (“P?”) or movie (“M?”) judgment for each image. The pleasantness judgment required participants to rate how pleasant they found the image (“Very”, “Moderate”, or “Not at all”) and the movie judgment required participants to determine which movie genre (“Action”, “Horror”, or “Comedy”) they believed was best associated with the object or scene. The response options for the cued judgment always appeared below the image. Responses for one judgment were entered with the right hand and responses for the other judgment were entered with the left hand. The hand assigned to each question was counterbalanced across participants. The instructions emphasized fast responses and that responding with the incorrect hand for a cued judgment counted as an incorrect response.

Participants were removed from the scanner following the study phase. After an approximately 15-minute break, participants began the self-paced recognition test which was completed outside the scanner in a behavioral testing room. Participants were shown the images from the study phase intermixed with images from the test phase and asked to indicate if they had studied the image (item recognition) and, if studied, what judgment (source memory) they made about the image during the study phase. This item and source judgment were combined into a single judgment with four options: “Old-Pleasant”, “Old-Movie”, “Old-Don’ t Know”, “New”. Participants were instructed to only respond with one of the three “Old” judgments if they had high confidence that the image was studied, and to respond “New” otherwise. Likewise, participants were instructed to only indicate they remember the source detail (i.e., “Old-Pleasant” or “Old-Movie”) if they had high confidence and to response “Old-Don’t Know” otherwise. Participants were instructed to enter responses quickly without sacrificing accuracy.

### Analysis of Memory Performance

For the present result, the analysis of the behavioral data focused on two measures of memory performance: item recognition and source memory. Item recognition was computed as the difference between the hit and false alarm rates for the initial old/new decision. Source memory was computed using a single-high threshold model (Snodgrass and Corwin, 1988) that accounts for the ‘ guess rate’ (e.g., Mattson et al., 2014):

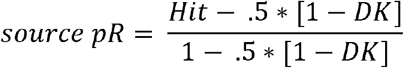

The *Hit* and *DK* variables in the above formula refer to the proportion of correct ‘ old’ responses accompanied by an accurate or don’ t know source memory judgments, respectively. Note that trials that received no response or a response with the incorrect hand during the study phase were excluded from the analysis.

### MRI Acquisition and Processing

MRI data were acquired at the University of Texas Southwestern Medical Center’ s Advanced Imaging Research Center with a 3T Philips Achieva MRI scanner (Philips Medical Systems, Andover, MA, USA) equipped with a 32-channel receiver head coil. Functional images were acquired with a blood oxygenation level dependent (BOLD) T2*-weighted echoplanar imaging (EPI) sequence (SENSE factor = 1.5, flip angle = 70°, 80 × 80 matrix, FOV = 240 mm x 240 mm, TR = 2000 ms, TE = 30 ms, 34 ascending slices, slice thickness = 3 mm, slice gap = 1 mm), and were oriented parallel to AC-PC. Five “dummy” scans were acquired at the start of each EPI session and discarded to allow for equilibration of tissue magnetization. A total of 180 functional volumes were acquired during each study session, for a total of 900 brain volumes. T1-weighted images (MPRAGE sequence, 240 × 240 matrix, 1 mm isotropic voxels) were acquired for anatomical reference prior to the first study session.

The functional data were preprocessed with Statistical Parametric Mapping (SPM12, Wellcome Department of Cognitive Neurology, London, UK) implemented in Matlab 2017b (The Mathworks, Inc., USA). The functional data were reoriented, subjected to a two-pass realignment procedure whereby images were first realigned to the first image of a session and then realigned to a mean EPI image, corrected for slice acquisition time differences using sinc interpolation with reference to the middle slice, and spatially normalized to a study specific EPI template (cf. de Chastelaine et al., 2011).

Trial-specific beta maps for each participant were obtained using a least-squares-all (LSA) general linear model (Mumford et al., 2014; Rissman et al., 2004). A regressor for each trial was modeled as a 2 s-duration boxcar convolved with a canonical hemodynamic response function. The fMRI data from the five study sessions were concatenated in time prior to the general linear model. Covariates of no interest in this first level model included the 6 rigid body motion parameters estimated from the realignment procedure and 4 session specific means (for sessions 2-5). Henceforth, trial-specific beta maps will be referred to as observed maps.

### Regions of Interest

The analyses reported here focus on the parahippocampal place area (PPA) and lateral occipital complex (LOC) regions of interest which are known to show selective increases in activation for scene (Epstein & Kanwisher, 1998) and object (Grill-Spector, 2003) stimuli, respectively. Briefly, bilateral PPA and LOC regions of interest were obtained from an unpublished dataset (N=22; 14 younger adults; 8 older adults) in which participants viewed images of scenes, faces, and objects in a blocked fMRI design. The PPA and LOC r (see Figure 1B) were identified from a second-level general linear model contrasting (at a voxel-wise threshold of *p* < .05, FWE-corrected) the BOLD response between scenes and objects (for full details of how these regions were defined, see Koen et al., 2019). The bilateral PPA and LOC were combined into a single region of interest for the present analyses. Note that the main conclusions were unchanged if the results described below were split by the PPA and LOC. The data broken down by ROI are provided on the Open Science Framework repository for this report.

### Across-Participant Pattern Similarity Analysis

#### Overview

The present analysis aimed to estimate stimulus (i.e., image) specific neural patterns using an across-participant pattern similarity analysis (Chen et al., 2017; Koch et al., 2020; Xiao et al., 2020; Zadbood et al., 2017; for related across-participant classification analyses, see Thakral et al., 2019). An overview of the procedure is depicted in Figure 1C-D. The present approach used linear mixed effects models to predict the estimated BOLD response (i.e., beta value) for each scene and object image in each voxel—henceforth referred to as predicted image maps. The advantage of using a linear mixed effects model is that the approach can statistically control for variables (e.g., image category, subsequent memory accuracy, etc.) that might reduce the ability to reliably measure stimulus specific neural activity from a single observed map of predicted beta responses from one participant. Moreover, this approach also allows for the estimation of predicted beta responses across all participants for these other factors, such as image category, which is also examined here to investigate age differences in category level similarity (see Introduction).

The analysis procedure used a leave-one-yoked-pair-out approach whereby data from the young and older adult pair with the same yoked stimulus set were held out and data from the remaining 23 young and 23 older adults were used to create the predicted image maps. The spatial patterns of the predicted image maps in the combined PPA and LOC of are then correlated with the observed image maps from the held out young and older adult. These correlations are used to compute both the item level and category level similarity measures. The following sections describe the analysis steps in more detail.

#### Estimation of Predicted Image and Predicted Category Maps

The steps for this part analysis are depicted in Figure 1C. For each iteration (out of 24 total), the trial-wise data from 23 young and 23 older adults was submitted to a linear mixed effects model to create the predicted image and category (i.e., scene and object) maps. A linear mixed effects model was created that predicted the beta response in a voxel with the fixed factors of participant age group (young vs. older), image category (scene vs. object), study judgment (pleasantness vs. movie), and subsequent memory (source correct, source incorrect, item miss; see Materials and Procedure). Note that these 4 variables were effect coded so that the global (fixed) intercept reflected the mean of all conditions. The model also included random intercepts on participant and image. No random slope components were included in the model. The equation for this linear mixed model can be written as:

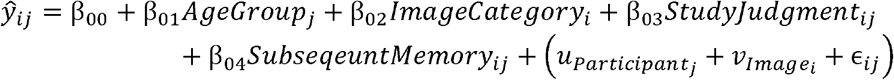

In the above equation, *ŷ*_*ij*_ is the predicted beta response for participant *j* and image *i*, 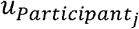 is the random intercept for participant *j*, and 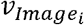 is the random intercept for image *i*. The *β*_s_ reflect the fixed effect coefficients for each of the predictor variables.

The predicted image maps were created from the random intercept term for image (i.e., 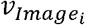) in each voxel. Importantly, this term in the model captures the predicted response (i.e., first-level beta value) in each voxel that is specific to a given image while controlling for variables that may also influence the estimated neural response in each voxel, such as image category. The predicted category maps were created from combining model coefficients for the intercept and image category terms (*β*_00_ − *β*_02_ for scenes; *β*_00_ + *β*_02_ for objects). The above procedure was carried out for all voxels in a grey matter mask defined by thresholding the grey matter tissue probability map in SPM12 (> .2 probability) with the intersection of each participant’ s implicit mask from the first level model. However, the primary analysis reported here focuses on the combined PPA and LOC region of interest (see Regions of Interest).

#### Computing the Item and Category Level Pattern Similarity Measures

Item and category level similarity measures were obtained for each studied image for each participant from a pattern similarity analysis conducted between the observed beta maps with the predicted image and category maps (see Estimation of Predicted Image and Predicted Category Maps; Figure 1D). The similarity metric for both measures was a Fisher *z*-transformed Pearson correlation coefficient (*r*_*z*_) and was based on features from all the voxels in the bilateral PPA and LOC regions of interest.

The computation of the item level similarity measure is depicted by the red lines in Figure 1D. The item level similarity measure required computing the pattern similarity between the observed and predicted maps for the same image and all images in the same category (i.e., within category). Specifically, the same image pattern similarity was computed as the *r*_*z*_ value between the observed beta map and the predicted image map for the same image (e.g., predicted image map for Scene 2 with observed beta map for Scene 2). The within category similarity measure was computed as the average *r*_*z*_ value between the observed beta map with the predicted image maps for all other images from the same category (e.g., observed beta map for Scene 2 with the predicted image map for all other scenes).

The category level similarity measure was computed as the difference between the observed beta map and the predicted category maps for scenes and objects (see blue lines and boxes in Figure 1D). This was scored such that positive values indicate higher levels of similarity between the observed beta map and predicted category map from the same category relative to the other category (i.e., within-minus-between category).

#### Simulation

A simulation was carried out to verify the across-participant pattern similarity method described above, and the results are reported in the Supplemental Material. Briefly, I generated data across 1000 ‘ voxels’ for 48 ‘ participants’ that mimicked the structure of the observed data with the addition of white noise. Half of the simulated participants were treated as young adults and half were treated as older adults, and yoked pairs of young and older adults were created. Note the simulation did not build in any age differences as the primary focus was to determine if the method can capture the item level and category level similarity measures. This simulated data was subjected to the above-described analysis method. The primary results from the stimulation demonstrated that the analysis captures both the item and category level similarity measures. For item level similarity, the *r*_*z*_ value for the same image similarity was greater than the *r*_*z*_ value for both the within and between category measures (see the red highlighted portions of Figure 1 C and D; for results, see Supplemental Figure 1). In relation to the item level similarity measure, it is worth noting that both the within and between category measures derived from the predicted image maps in the simulated data were no different than 0, which is consistent with the findings from the findings from the observed data reported below (see Supplemental Figure 2). For the category level similarity measure, the *r*_*z*_ value was higher with the within category map (e.g., observed beta map for a scene with the predicted scene map) than with the between category map for the simulated data. This is also consistent with the observed data for the category level similarity measure (see Supplemental Figure 3 and 4). Overall, the simulation results suggest that the across-participant pattern similarity analysis can be used to measure both item and category level pattern similarity.

### Statistical Analyses

All statistical analyses were carried out in R (R Core Team, 2021). A significance threshold of *p* < .05 was used for all analyses unless otherwise indicated.

#### Neuropsychological Test and Memory Performance

Independent samples *t*-tests (assuming unequal variances) were used to examine age differences in performance on the neuropsychological test battery. The item and source memory measures were submitted to separate 2 (Age Group: Young, Older) X 2 (Image Type: Object, Scene) mixed-factorial ANOVA as implemented in the *aov_ez* function of the *afex* package (Singmann et al., 2021).

#### Item and Category level Pattern Similarity Measures

Statistical analyses of both item and category level similarity metrics were carried out using linear mixed effects models as implemented in the *mixed* function of the *afex* package. For clarity, these models are essentially an analysis of covariance as implemented in a linear mixed effects model. Specifically, this analysis included three factors (age group, image category, subsequent memory) and their interactions.

Note that the subsequent memory factor had three levels: source correct (item hits with a correct source judgment), source incorrect (item hits with an incorrect or a don’ t know source judgment), and item misses (incorrect new response) to a studied image. For the analysis of item level similarity, the category level similarity measure was included as a covariate. Likewise, item level similarity was included as a covariate for the category level similarity analysis. Note that both models included random intercepts (or error terms) for participant and image. Significance of the factors of interest (age group, image category, subsequent memory, and their interactions) were assessed using likelihood ratio tests. The average number of trials in each of the cells formed by the age group, image category, and subsequent memory factors are reported in Supplemental Table 2. Post-hoc tests were conducted with the *emmeans* (Lenth, 2021) package using asymptotic degrees of freedom (i.e., *z* tests). Additionally, where appropriate, post-hoc tests were corrected for multiple comparisons using the Holm procedure (Holm, 1979). Note that the results for the analysis of item and category level similarity measures were unchanged when computing degrees of freedom with the Satterthwaite (1946) method.

#### Analysis of Inter-individual Differences

The item and category level similarity measures were also related to memory performance on the experimental recognition memory and a univariate differentiation index reported in the previous analysis of this dataset (Koen et al., 2019). For these analyses, a single item and category similarity measure was computed for each participant by averaging the measure across all studied trials. Briefly, the differentiation index is calculated as the difference between the magnitude of the BOLD response between a region’ s preferred and not preferred stimulus category scaled by a pooled standard deviation (akin to a discrimination index from signal detection theory). Here, the differentiation index was averaged across the PPA and LOC to mimic the procedure for computing the across-participant derived item and category level similarity measures, which were computed from a combined PPA and LOC region of interest. The memory measures included item recognition and source memory (see Analysis of Memory Performance). Both memory measures were averaged across the image category to get a single measure of memory performance per participant. Associations between the above measures were examined using partial correlations. Linear regressions were also conducted to test if the associations were moderated by age group.

## Results

### Neuropsychological and Memory Measures

Performance on the neuropsychological test battery revealed a standard pattern of age differences in cognition (see Supplemental Table 1). Older adults showed lower performance compared to younger adults on tests of declarative memory, processing speed, category fluency, and reasoning, but there were no age differences in measures of verbal fluency and reading.

The item and source memory measures are show in in Table 1. The 2 (age group) X 2 (image Type) mixed factorial ANOVA on the item recognition measure revealed a significant main effect of image type, *F*(1,46) = 187.97, *MS*_e_ = .01, *p* < 10^−15^, partial-*η*^2^ = .80, reflecting better item recognition for objects than scenes. The main effect of age group, *F*(1,46) = 3.89, *MS*_e_ = .04, *p* = .055, partial-*η*^2^ = .08, and the interaction between age and image type, *F*(1,46) = 1.04, *MS*_e_ = .01, *p* = .312, partial-*η*^2^ = .02, were not significant. The mixed factorial ANOVA on the source memory also produced a significant main effect of image type, *F*(1,46) = 105.05, *MS*_e_ = .01, *p* < 10^−12^, partial-*η*^2^ = .70, which was driven by better source memory for objects than scenes. Likewise, there was no significant main effect of age group, *F*(1,46) = .81, *MS*_e_ = .06, *p* = .372, partial-*η*^2^ = .02, nor a significant interaction between age group and image type, *F*(1,46) = .97, *MS*_e_ = .01, *p* = .329, partial-*η*^2^ = .02. In summary, although a standard pattern of age differences was observed on the neuropsychological test battery, there were no reliable age differences observed on the experimental memory task.

**Table 1.**
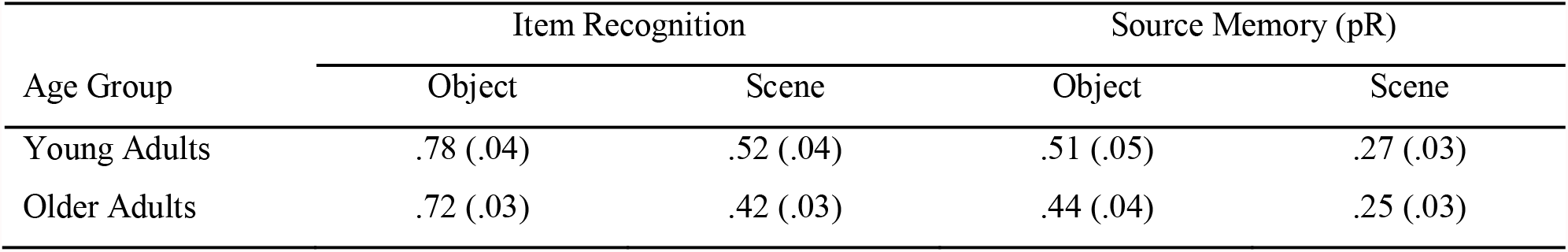
Means and standard errors for the item recognition and source memory measures.

| Age Group | Item Recognition |  | Source Memory (pR) |  |
| --- | --- | --- | --- | --- |
|  | Object | Scene | Object | Scene |
| Young Adults | .78 (.04) | .52 (.04) | .51 (.05) | .27 (.03) |
| Older Adults | .72 (.03) | .42 (.03) | .44 (.04) | .25 (.03) |

### Item level Pattern Similarity

The linear mixed effects for item level similarity revealed two significant effects (for a description of the model, see Statistical Analyses). First, there was a main effect of age group [*χ*^2^(1) = 6.35, *p* = .012] that was driven by item level similarity being reduced in older adults compared to younger adults (see Figure 2A). This finding is consistent with the prediction that increasing age is associated with dedifferentiation of item specific representations. Importantly, both young adults [*z* = 7.42, *p* < .001] and older adults [*z* = 5.363, *p* < .001] showed item similarity measures significantly greater than 0, indicating that item level similarity was reduced, but not entirely absent, in older adults. Importantly, this age difference was driven by differences in the same image component of the item similarity measure (see Supplemental Figure S2 for the *r*_*z*_ values for the three component scores derived from the predicted image maps).

Second, and importantly, there was a significant interaction between age group and subsequent memory [*χ*^2^(2) = 9.20, *p* = .011] (see Figure 2B). To examine this interaction, I first conducted a post-hoc test that examined effects of subsequent memory within young and older adults separately. Young adults showed significantly higher item level similarity for source correct trials (i.e., correct recognition of the image and encoding task) relative to item misses[*z* = 3.170, *p* = .002, *p*_Holm_ = .01], along with no differences between source correct and incorrect trials (i.e., correct recognition of the image but failure to correctly recognize the encoding task) [*z* = 1.486, *p* = .137, *p*_Holm_ = .411], or between source incorrect and item miss trials [*z* = 1.789, *p* = .074, *p*_Holm_ = .368]. This pattern suggests that there was a graded positive subsequent memory effect in young adults. In contrast, older adults showed no reliable effects of subsequent memory on measures of item level similarity [source correct vs. item miss: *z* = −0.802, *p* = .422, *p*_Holm_ = .479; source correct vs. incorrect: *z* = 1.176, *p* = .239, *p*_Holm_ = .479; source incorrect vs. item miss: *z* = −1.756, *p* = .079, *p*_Holm_ = .368]. Second, I decomposed this age group by subsequent memory interaction by contrasting young and older adults within each level of subsequent memory. This analysis revealed that young adults had significantly higher levels of item level similarity relative to older adults for source correct [*z* = 3.532, *p* < .001, *p*_Holm_ = .001] and source incorrect trials [*z* = 2.849, *p* = .004, *p*_Holm_ = .009], but not item misses, [*z* = −0.473, *p* = .636, *p*_Holm_ = .636].

**Figure 2.**
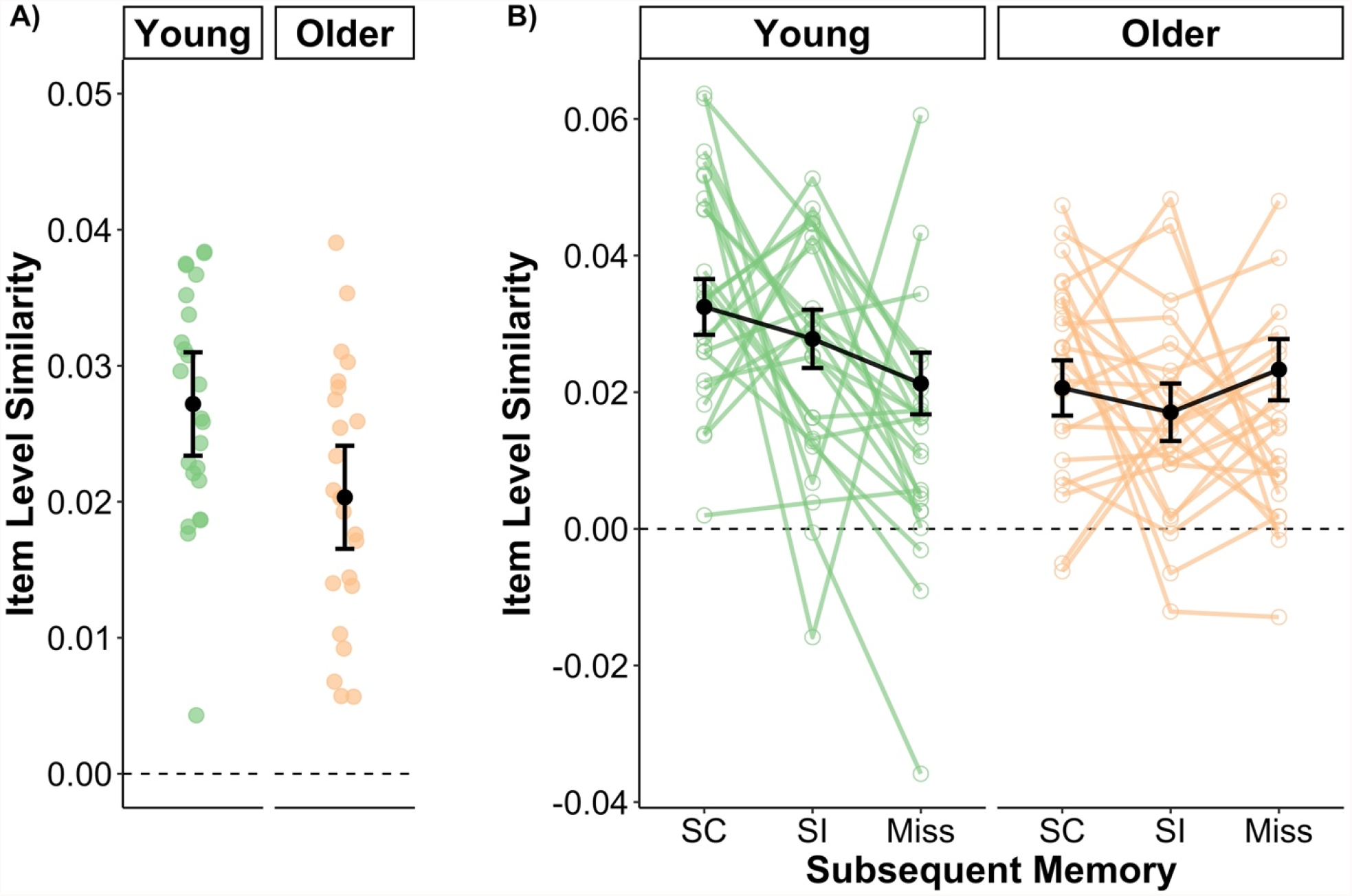
Measures of item level similarity for young and older adults and their relationship to subsequent memory. (A) Item level similarity for young adults was significantly higher than that of older adults suggesting that age is associated with dedifferentiation of item-specific neural representations. (B) Item level similarity differed as a function of subsequent memory in young, but not older adults. The effect in young adults was graded such that item level similarity was highest during trials that attracted correct item and source memory decisions (SC) and lowest for trials in which the image was subsequently forgotten. The observed (green and orange) points reflect the unadjusted mean category similarity measure across all images and other factors not included in the plots for each participant. Note that the black points represent the model implied group mean and error bars (black points) controlling for other terms in the model, including category level similarity. Error bars represent ± 1 standard error of the mean.

In summary, the above results show that item level similarity in young adults, but not in older adults, was modulated by subsequent memory in a graded fashion. Moreover, these results are consistent with predictions that age is associated with a reduction in the fidelity of neural representations for individual items.

### Category level Pattern Similarity

The linear mixed effects model for category level similarity revealed a main effect of age group [*χ*^2^(1) = 25.234, *p* < .001] that was driven by higher category level similarity in young adults relative to older adults. Note that this age difference appears to be driven by reductions in within-category component of the category level similarity measure, but not the between-category component (see Supplemental Figure 4). There was also a main effect of image category reflecting high category similarity for scenes relative to objects [*χ*^2^(1) = 23.903, *p* < .001]. These two main effects were qualified by a significant age group by image category interaction [*χ*^2^(2) = 53.022, *p* < .001]. Post-hoc tests of this age group by image category interaction revealed that age differences were larger for scenes [*z* = 8.172, *p* < .001] than for objects [*z* = 2.601, *p* = .009] (Figure 3A). This pattern of results is consistent with our prior analysis of this dataset showing that age differences in a univariate measure of neural differentiation were significant for scenes but not objects (Koen et al., 2019; see also, Srokova et al., 2020).

**Figure 3.**
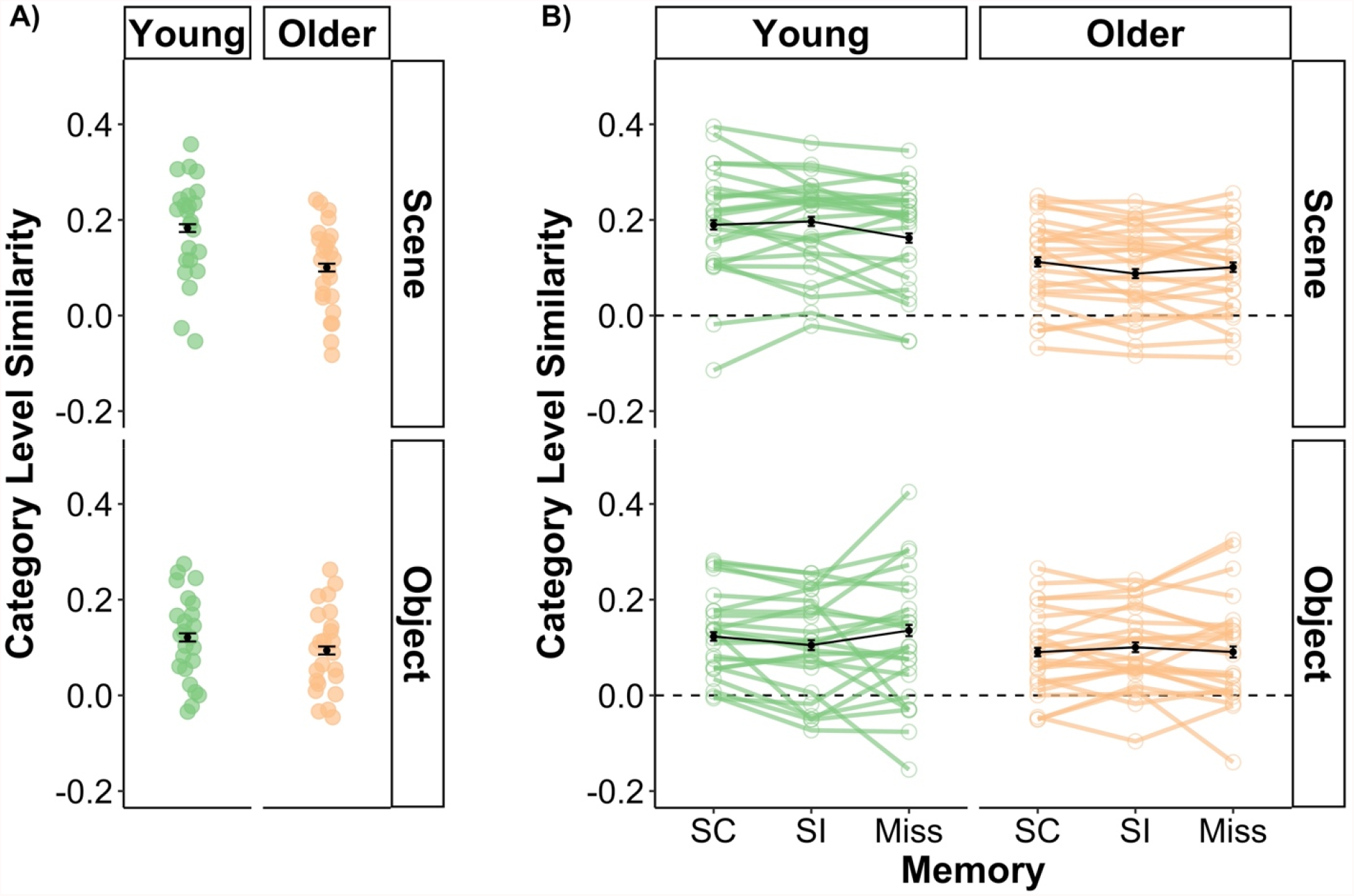
(A) Measures of category level similarity for young and older separated by objects and scenes. Category level similarity was significantly higher in young compared to older adults for both scenes and objects. However, as evident in the figure, this age difference was more robust for scene images. (B) Visual depiction of the three-way interaction involving age, image category, and subsequent memory for measures of category level similarity. The interaction was driven by age differences in the pattern of subsequent memory effects for scenes, but not objects. Young adults demonstrated higher category level similarity for source correct (SC) and source incorrect (SI) trials relative to item misses. In contrast, older adults only showed elevated category level similarity between SC and SI scene trials. For objects, there were no significant subsequent memory effects in either age group. The observed (green and orange) points reflect the unadjusted mean category similarity measure across all images and other factors not included in the plots for each participant. Note that the black points represent the model implied group mean and error bars (black points) controlling for other terms in the model, including category level similarity. Error bars represent ± 1 standard error of the mean.

There were also two interactions involving subsequent memory: a category by subsequent memory interaction [*χ*^2^(2) = 7.377, *p* = .025] and an interaction between age group, image category, and subsequent memory [*χ*^2^(2) = 21.625, *p* < .001]. Here, I focus on decomposing the three-way interaction (see Figure 3B). This interaction was driven by age differences in patterns of subsequent memory effects for scenes. Note that post-hoc tests showed no age or subsequent memory differences were present for objects [all *p*’ s_Holm_ > .075]. The post-hoc tests reported below are specific to scene stimuli. In young adults, category level similarity for scenes was higher for source correct [*z* = 2.945, *p* = .003, *p*_Holm_ = .035] and source incorrect trials [*z* = 3.886, *p* < .001, *p*_Holm_ = .001] relative to item misses, with no significant difference between source correct and incorrect trials [*z* = −0.810, *p* =.418, *p*_Holm_ = 1.00]. Put differently, category level similarity for scenes in young adults tracked subsequent item recognition, regardless of source memory accuracy (cf. Mattson et al., 2014; Ranganath et al., 2004). However, in older adults, category level similarity for scenes was only significantly higher for source correct relative to source incorrect trials [*z* = 2.856, *p* = .004, *p*_Holm_ = .043]. No significant differences were observed between source correct and item miss trials [*z* = 1.213, *p* = .225, *p*_Holm_ = 1.00] nor between source incorrect and item miss trials [*z* = −1.49, *p* =.136, *p*_Holm_ = .953]. It is important to note that, while not significant, source correct trials had numerically higher category level similarity compared to item misses. For older adults, the pattern of subsequent memory effects appeared to track source memory accuracy but not item recognition.

### Association with Univariate Differentiation Index

The next analysis used partial correlations to examine the inter-individual relationships between the item level and category level similarity measures presented above and the univariate differentiation index from the initial analysis of this dataset (Koen et al., 2019). Briefly, the univariate differentiation index is the difference between a regions mean response to its preferred and non-preferred stimulus category scaled by a pooled standard deviation (cf. Koen et al., 2019; Srokova et al., 2020; Voss et al., 2008; Zebrowitz et al., 2016). Importantly, the differentiation index is thought to capture neural distinctiveness for category, but not item level, information. Thus, it reasons that the univariate differentiation index will be correlated with the category level similarity measure, but not the item level similarity measure, given that they are both hypothesized to capture the same level of information.

For this analysis, a single univariate differentiation index was computed by averaging the differentiation indices for the PPA and LOC to mimic how the item and category level similarity measures were computed in the present report (for additional details, see Koen et al., 2019). Likewise, for each participant, a single measure of item level and category level similarity was obtained by averaging across all trials. Note that the zero-order correlation between the item and category level similarity measures was significant [*r*(46) = .725, *p* < .001]. Thus, like the linear mixed effects models reported on previously, the partial correlation between item level similarity and the univariate differentiation index controlled for category level similarity (in addition to age group), and vice versa for the partial correlation involving category level similarity and the univariate differentiation index.

The results of this analysis are shown in Figure 4A and 4D. The partial correlation between the univariate differentiation index and the category level similarity measure (controlling for age group and item level similarity) was significant [*r*_partial_(44) = .691, *p* < .001]. In contrast, the partial correlation between the univariate differentiation index and item level similarity (controlling for age group and category level similarity) was not significant [*r*_partial_(44) = −.083, *p* = .584]. Note that a multiple linear regression analysis provided little support to suggest that age moderated the relationship between item and category level similarity with the univariate differentiation index [*p*’ s > .352]. In summary, these findings show that the category level similarity measure, but not the item similarity measure, derived from the across-participant approach is most strongly associated with the univariate differentiation index. This suggests that the category level similarity measure and univariate differentiation index likely measure a similar underlying process but may be differentially sensitive to intra-and inter-individual differences in memory.

**Figure 4.**
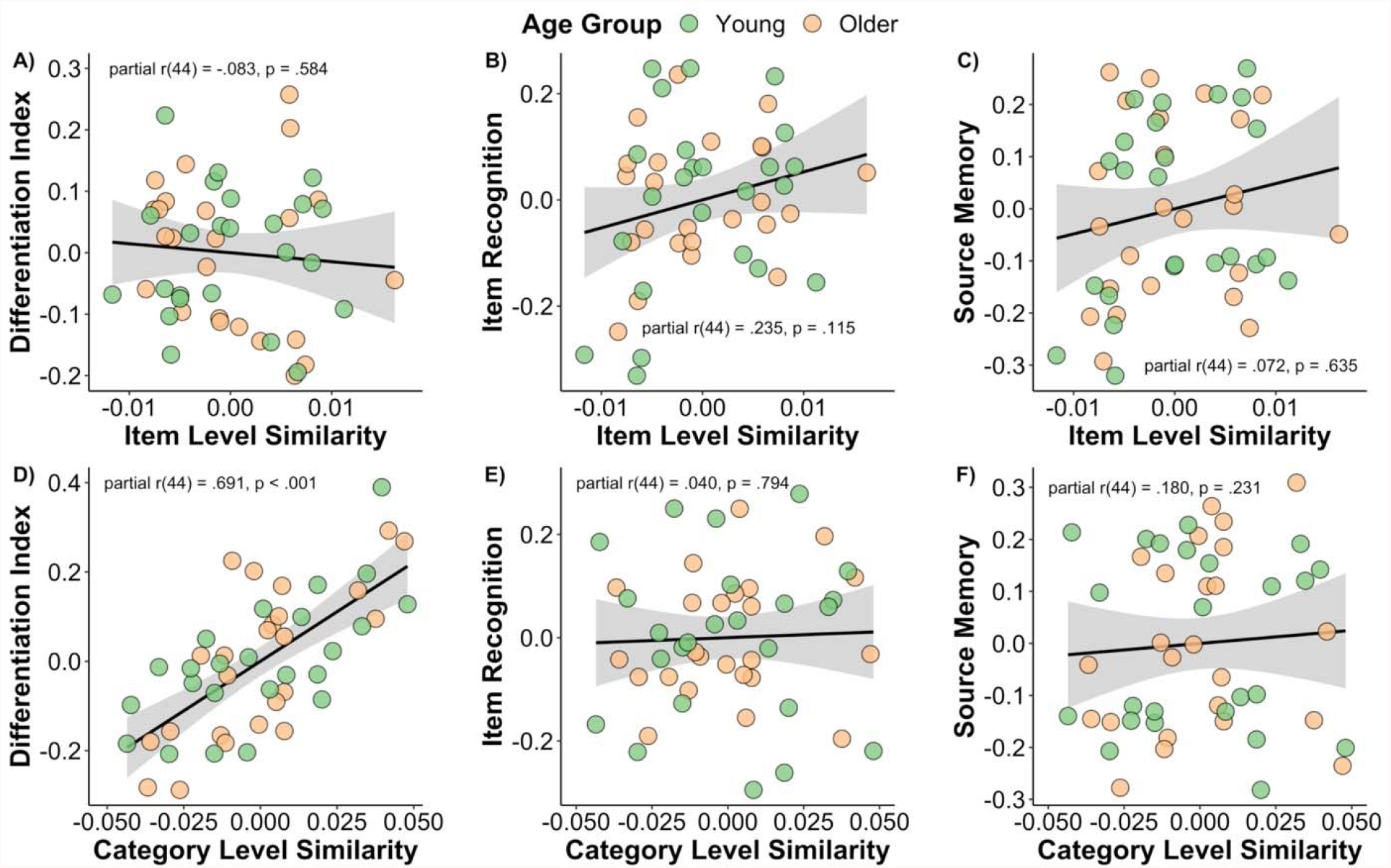
Partial correlation plots depicting the relationships between measures of item and category level similarity with univariate measures of neural differentiation (i.e., differentiation index) and two measures of recognition memory performance. The only significant relationship observed was between the category level similarity measure and the univariate differentiation index for this data initially reported by Koen and colleagues (2019). Note that plots in panels A-C control for age group and category level similarity, whereas the plots in panels D-F control for age group and item level similarity.

### Associations with Item Recognition and Source Memory

A series of partial correlations were also conducted to examine the relationship between the similarity metrics (i.e., item and category level) with individual differences in memory performance (i.e., item recognition and source memory). The two partial correlations (controlling for age group and category level similarity) between item level similarity and memory performance were not significant [item recognition: *r*_partial_(44) = .235, *p* = .115; source memory: *r*_partial_(44) = .180, *p* = .231] (Figure 4B and 4C). Similarly, the two partial correlations (controlling for age group and item level similarity) involving category level similarity and memory were not significant [item recognition: *r*_partial_(44) = .049, *p* = .794; source memory: *r*_partial_(44) = .071, *p* = .635] (Figure 4E and 4F). In short, there was little indication that the item and category level similarity measures explained inter-individual differences in memory performance.

### General Discussion

The present study investigated age differences in item and category level neural representations in the ventral visual cortex (specifically in the PPA and LOC) and how these different levels of neural representations were related to successful memory encoding. Here, I applied a across-participant pattern similarity analysis to a previously published dataset (Koen et al., 2019) to measure patterns of neural activity specific to individual stimuli (item level similarity) and neural patterns for categories of visual stimuli (e.g., scenes and objects; see Figure 1). Across-participant pattern similarity analyses can capture stimulus specific patterns of neural activity (Chen et al., 2017; Koch et al., 2020; Xiao et al., 2020; Zadbood et al., 2017) as well as patterns of activity for categorical features (Thakral et al., 2019), making it an strong approach to simultaneously investigate age-related dedifferentiation across different levels of neural representation. There were four main findings. First, older adults showed a significant reduction in both measures of item and category similarity relative to young adults (Figure 2A). Second, item level similarity showed a graded, positive subsequent memory effect in young adults, but no subsequent memory effect in older adults (Figure 2B). In contrast, category level similarity for scene stimuli showed subsequent memory effects consistent with item recognition in young adults and subsequent source memory, but not item recognition, in older adults (see Figure 4). Lastly, category, but not item level, similarity correlated with a univariate measures of category selectivity (i.e., the differentiation index; cf. Koen et al., 2019; Voss et al., 2008). Below, I discuss these findings and the across-participant pattern similarity approach used in this study.

### Behavioral Findings

One unexpected finding was the lack of significant age differences in measures of both item and source memory on the experimental memory task (for a previous discussion on this issue with this participant sample, see Koen et al., 2019; also see, Horne et al., 2020; Koen et al., 2018). The lack of a significant difference in source memory is particularly surprising given the robust literature on large age-related reductions in recollection (Koen & Yonelinas, 2014; Old & Naveh-Benjamin, 2008; Schoemaker et al., 2014; Spencer & Raz, 1995). While there are many possibilities for this null finding, one that stands out is that the encoding task might have benefited successful memory encoding processes in older adults more than young adults, which has previously been shown to eliminate age differences in recollection (Luo & Craik, 2008). Another related possibility is that there were multiple redundant pieces of information that can support success retrieval (see Horne et al., 2020). For a participant in the present task, a given study judgement (e.g., pleasantness judgment) was consistently paired with a unique cue (e.g., ‘ P?’) and a specific response hand (e.g., right hand). These multiple redundant features during study may have differentially benefitted older adults’ memory be increasing the efficacy of successful encoding processing or providing more routes to retrieve task relevant information. Future research should test if using redundant versus overlapping features during encoding differentially affects young and older adults’ memory performance.

### Item and Category Level Similarity

Turning to the neuroimaging findings, the results reported here support a key prediction of computational models of neural dedifferentiation (Li et al., 2001; Li & Sikström, 2002; Li & Rieckmann, 2014). These models argue that neural dedifferentiation results from age-related decreases in neuromodulatory drive that, in turn, reduce the distinctiveness of neural representations for individual stimuli (i.e., item level representations) and broad categories of stimuli (i.e., category level representations). The results also converge with recent fMRI findings showing age-related dedifferentiation for different levels of neural representations (Kobelt et al., 2021; Trelle et al., 2019; for review, see Koen & Rugg, 2019; Koen et al., 2020). In addition, the current findings extend this prior research by showing that across-participant pattern similarity analyses are sensitive to age differences across both item and category level similarity measures. It is important to point out that the present results were driven by a reduction, not an absence, of item and category level similarity in older adults. Although item and category level similarity were highly correlated, age differences in item level similarity were independent of age differences in category level similarity (and vice versa). This suggests that age-related neural dedifferentiation for different levels of neural representations might be driven by potentially independent mechanisms that, in turn, have different contributes to memory and cognitive aging.

The present findings also suggest that the neural representations that support successful memory encoding differ between young and older adults. This is supported by observing age differences in the patterns of subsequent memory effects for item and category level similarity. In young adults, item level similarity showed a graded positive subsequent memory effect in which item level similarity was highest for trials that were recognized and had the source memory detail (i.e., the encoding task) remembered, and lowest for trials that went on to be forgotten. This suggests that item level similarity is linearly related to the amount of information that is accessible on a subsequent memory test in young adults (e.g., Kobelt et al., 2021; Koch et al., 2020; Zheng et al., 2018). In contrast, item level similarity was not modulated by subsequent memory in older adults. Moreover, the age differences in item level similarity were largest for trials in which the source was accurately remembered (i.e., source correct trials) whereas no age differences were observed for trials that went on to be forgotten. One interpretation of the age group by memory interaction described above is that older adults do not form neural representations of events during encoding as distinct as their younger counterparts do (cf. Craik & Rose, 2012; Friedman & Johnson, 2014). Another possibility is that age-related reductions in item level similarity are driven by older adults having less across-participant consistency in neural representations, and not a decline in the representational distinctiveness *per se*. Differences in across-group consistency can presumably arise for multiple reasons, such as age differences in life experience (Koen & Rugg, 2019) or differences in neural noise (Li et al., 2001; Voytek et al., 2015), and the reason for this may have important implications for how neural dedifferentiation influences memory. For instance, finding that increased neural noise results in age-related decreases in across-participant consistency would align well with computational models of neural dedifferentiation which argue that neural noise, resulting from age-related decreases in neuromodulatory drive, result in neural dedifferentiation. Future research investigating these issues is warranted.

Unlike item level similarity, measures of category similarity showed significant subsequent memory effects in both young and older adults, albeit only for scene stimuli. In young adults, the effect appeared to be related to encoding processes supporting subsequent successful item memory, but not source memory. In contrast, older adults showed a pattern of subsequent memory effects for category level similarity whereby only source correct trials were greater than source incorrect trials, but not item miss trials. This finding does not easily fit with supporting item or source memory. Nonetheless, the present findings demonstrate that a full understanding of the role of neural dedifferentiation to cognitive aging will benefit from investigations that simultaneously quantifies dedifferentiation across different levels of representation.

Interpreting the above age differences in both item and category level similarity must consider the lack of age differences in the measures of item and source memory. First, the present results only apply to the combined bilateral PPA and LOC region of interest used in the present analysis. Thus, it is possible that the item and category level similarity may show more robust subsequent memory effects in regions not considered here, such as the frontal cortex or more anterior regions of temporal cortex (Deng et al., 2021). For example, Deng and colleagues (2021) recently demonstrated that, whereas there was age-related reductions in the distinctiveness of patterns in visual cortex, there were age-related increases in the distinctiveness of neural patterns for conceptual and mnemonic representations in the anterior temporal lobe, a finding they termed hyperdifferentiation. A second possibility is that, in addition to deficits at encoding, older adults also have greater variability in the engagement of retrieval processes that support successful memory. It could be that successful retrieval processes are more closely aligned with successful encoding processes in young adults, but not older adults. This additional variance at retrieval in older adults, which was not accounted for in the present study, could help explain why we observed differences in the neural data (i.e., item and category level similarity) but not in the behavioral data. Note that the above possibilities are not mutually exclusive. Either, or both, of these factors can contribute to the observed age-related reductions in item and category level similarity in the face of no significant age differences in memory performance on the experimental task^1^.

### Brain-Behavior Relationships

The present analysis, however, did not detect any significant relationship between the across-participant item and category similarity measures with inter-individual differences in memory performance. These null findings conflict with prior research demonstrating that measures of neural selectivity (or distinctiveness) for both the item (Kobelt et al., 2021; Yassa et al., 2011) and category level (Berron et al., 2018; Koen et al., 2019; Srokova et al., 2020) correlate with inter-individual differences in memory performance. The null brain-behavior correlation was somewhat more surprising for the category level similarity measure given the strong correlation between this measure and the univariate differentiation index which, in the initial report of this dataset, correlated with item recognition memory performance (Koen et al., 2019). The reason for the discrepancy is unclear and should be addressed with future research. First, although the two neural measures (i.e., the multivariate category level similarity and univariate differentiation index) are highly correlated, the small sample size of the current study may have been underpowered to detect any relationship, should it exist, with the aggregate across-participant similarity measures. An alternative possibility is that the discrepancy resulted from the across-participant pattern similarity analyses is insensitive to individual differences in memory and cognition because they eliminate, or average out, the idiosyncratic patterns specific to a participant. It could be these participant-specific neural patterns, which within-participant methods are better able to capture, are important for revealing brain-behavior relationships in memory. A third possibility is that item level similarity was controlled for in the present study but not in the prior report of this data as it was not available at that time (Koen et al., 2019). Nonetheless, the results discussed above suggest that across-versus within-participant derived measures of category level representations might be differentially sensitive to intra- and inter-individua differences in memory.

### Implications for Across-Participant Pattern Similarity Multivariate Methods

The results reported here also have implications for the growing number of studies using across-participant pattern similarity analyses to investigate the neural underpinnings of memory and cognition (Chen et al., 2017; Koch et al., 2020; Xiao et al., 2020; Zadbood et al., 2017). Unlike traditional univariate methods, across-participant pattern analyses can potentially uncover patterns of neural activity that are specific to individual stimuli or events (but see Davis et al., 2014). Across-participant multivariate approaches also offer potential advantages over the more common within-participant approaches, specifically for measuring stimulus-specific neural patterns for experimental tasks that use a single presentation of a stimulus. As noted in the Introduction, within-participant pattern similarity approaches to measuring stimulus specific neural patterns rely on comparing the spatial profile of neural activation for repetitions of the exact stimulus (or a highly similar stimulus). Measures obtained from this approach can change across repetitions (cf. St-Laurent & Buchsbaum, 2019) which might influence comparisons between groups or conditions. Across-participant pattern similarity analyses can be applied to paradigms in which participants are shown each stimulus on a single occasion, thus obviating issues caused by within-participant stimulus repetition.

It should also be pointed out that the implementation of across-participant pattern similarity analysis differed from prior research which only correlated observed patterns of neural activity between participants (Chen et al., 2017; Koch et al., 2020; Xiao et al., 2020; Zadbood et al., 2017). Here, I used an iterative approach whereby linear mixed models were used to generate predicted patterns or neural activity using data from a ‘ training set’. These predicted maps were then correlated with observed maps of neural activity in participants held out of the training set (i.e., the ‘ test set’) to generate item and category level similarity measures. While both approaches are similar in their aims, I believe there are two features of the method developed here that might be particularly beneficial. First, the assumption of across-participant pattern similarity analyses is that this approach can identify the pattern of neural activity specific to an individual stimulus. This was achieved in prior studies by averaging the similarity measure obtained from correlating a single participant’ s neural pattern for a given stimulus with the pattern from all other participants who experienced the same stimulus. While this is a sound approach, a potential advantage of using linear mixed effects models is that one can more explicitly model, and in theory quantify, pattern of neural activity for a given stimulus (this is the random intercept term for the stimulus images in the present study). Moreover, this approach allows one to statistically control for factors (such as subsequent memory condition or age group) that might confound pattern similarity measures which might also help to quantify stimulus (or category specific) patterns more precisely. The second advantage of the linear mixed modelling approach used here is that it pools across trials and participants versus computing similarity measures across participants that have trials in a specific condition (e.g., remembered versus forgot a trial) to estimate the stimulus specific neural patterns. These two features, together, might potentially increase the sensitivity of across-participant pattern similarity analyses which can be beneficial to studies with small sample sizes, such as that in the present study. One potential limitation of the approach proposed here is that misspecification of the model used to create the predicted brain maps might reduce the across-participant derived similarity measures. It should be noted that this is one explanation of the lower item and category level similarity measures obtained in the observed data relative to the simulation (see Supplemental Material). While additional research is needed to better understand the benefits and limitations of across-pattern multivariate analyses (regardless of the specific implementations), the present findings add to the existing evidence that across-participant approaches are a promising avenue to advance our understanding of the neural underpinnings of memory and cognition across the lifespan.

### Conclusions

In conclusion, the present findings join a growing body of work showing that age-related neural dedifferentiation occurs for item and category level neural representations. Across-participant pattern similarity analyses revealed that older adults showed lower estimates of both item and category level similarity. Additionally, the item and category similarity measured during a memory encoding phase were differentially associated with successful subsequent memory in young and older adults. These findings suggest that neural dedifferentiation for item and category level neural representations might be associated with different aspects of age and memory. Future research will benefit from understanding the extent to which neural dedifferentiation across multiple representational levels, including those for large-scale neural networks (Cassady et al., 2019; Chan et al., 2014, 2017; for reviews, see Damoiseaux, 2017; Goh, 2011; Koen et al., 2020), are related to one another and influence patterns of cognitive aging.

## Acknowledgements

This work was supported by awards from the National Institute on Aging [grant numbers F32AG049583 and R56AG068149] and the Aging Mind Foundation to Joshua D. Koen. I would like to thank Dr. Michael Rugg for allowing me to reanalyze data collected as part of an award from the National Institute on Aging [grant number R01AG039103] for the present report. I would also like to thank Rachelle Pichot for providing feedback on an initial draft of this manuscript.

## Footnotes

1 I would like to thank an anonymous reviewer for recommending these alternative accounts.

