## Supplemental Material for "Age-related neural dedifferentiation for individual stimuli: An across-participant pattern similarity analysis"

Joshua D. Koen^1*^

^1^University of Notre Dame, Notre Dame, IN

**Data Availability Statement:** The data contributing to the analyses reported here are available on the Open Science Framework (<https://osf.io/hbuq9/>).

### Simulation of The Across-Participant Pattern Similarity Analysis

#### Method

I conducted a simulation to verify that the proposed across-participant pattern similarity analysis method in the main text could recover item and category level information. For the sim simulation, I generated data patterns over 1000 ‘voxels’. The simulated data were generated using the same linear mixed model used in Step 1 of the analysis, namely:

$$\hat{y}_{ij}=\beta_{00}+\beta_{01}AgeGroup_{j}+\beta_{02}ImageCategory_{i}+\beta_{03}StudyJudgment_{ij}+\beta_{04}SubseqeuntMemory_{ij}+\left( u_{Participant_{j}}+v_{Image_{i}}+\epsilon_{ij} \right)$$

In the above equation, $\hat{y}_{ij}$ is the predicted beta response for participant *j* and image *i*, $u_{Participant_{j}}$ is the random intercept for participant *j*, and $v_{Image_{i}}$ is the random intercept for image *i*. The $\beta$’s reflect the fixed effect coefficients for each of the predictor variables.

To simulate the data, a ‘beta’ map was created for each fixed effect term in the model. This included maps for the intercept, young adults, older adults, scene images, object images, pleasantness judgment, move judgment, subsequent source hits, subsequent source misses, and subsequent item misses. Additionally, image maps were created for 240 images (120 objects and 120 scenes) and for 48 participants (24 young and 24 older adults). The images were randomly assigned to the subsequent memory bins to mimic the distribution observed in the real data (collapsed over age because there was no significant difference in memory with respect to age). Yoked pairs of one young and one older adult were also created to mimic the analysis in the main text.

The voxel maps for fixed and random effects betas were generated from multivariate normal random distribution. These beta maps were added together, along with random white noise, to form an image map for each participant and image that served as the ‘observed’ beta map (see Supplemental Table 2 below). Note that correlations between levels of factors (e.g., age group, image, image category) were built into to the simulation based on the assumption that there would be at least some overlap in the patterns of neural activity between levels of the same factor.

The simulated data were the subjected to the analysis pipeline outlined in the main text (see Across-Participant Pattern Similarity Analysis and Figure 1 in the main text). For each of 24 yoked pairs, one yoked pair was held out for the test set and the data from the remaining 23 young pairs (1 young and 1 older adult each for a total of 46 participants) were used for the training set. Once the predicted image maps were created from the 23 young and 23 older simulated datasets used in training, the held out young and older adult simulated datasets were used to compute the image and category level similarity measures.

The focus on the simulation is on the component values (i.e., Fisher transformed Pearson *r* values, or $r_{z}$) used to compute the item and category level similarity measures. For the image similarity measure, the $r_{z}$ values for the same item, within category, and between category measures from the predicted image maps (i.e., that from the $v_{Image_{i}}$ term) are plotted. If the image specific similarity can be captured by this measure, then we should observe that the $r_{z}$ for the same image is greater than the $r_{z}$ for both the within and between category predicted image maps. Note that a comparison of the same item and within category $r_{z}$ in this manner is equivalent to the same item - within category measure difference score used in the main. For the category level similarity measure, we expect that the observed images with show a stronger $r_{z}$ with the within category map compared to the between category map. This comparison of the within and between category $r_{z}$ values (from the predicted category maps) is equivalent to the within-minus-between category similarity measure used in the main text.

#### Results

##### Item Level Similarity

The results from the simulation for the item level similarity measure are shown in Supplemental Figure 1. This figure presents the $r_{z}$ values for each participant (averaged across images) for the three component measures from computing the observed beta map with the predicted image map (i.e., same image, within category, and between category). As can be seen in Supplemental Figure 1, the same image $r_{z}$ is non-zero whereas the within and between category $r_{z}$ hover around zero with very little variance. This pattern indicates that the proposed across-participant analysis method can capture item (or image) specific patterns of neural activity. When compared to Supplemental Figure 2, which plots the same measures from the observed data. One thing to note is that while the $r_{z}$ measure for the within and between category predicted image maps are comparable (i.e., both near 0), the same image $r_{z}$ is higher for the simulated data relative to the same image $r_{z}$ obtained from the observed data. These differences are expected to some extent between observed and simulated data. This can be due to higher noise in the observed data than in the simulated data, omission of model terms used in the linear mixed model used to create the predicted maps in the observed data (e.g., interactions between factors or random slopes), or how noise was introduced in the simulation (e.g., white versus pink noise). Critically, however, the pattern across the simulated and observed data is similar with the component $r_{z}$ measures that comprised the item level similarity measure (i.e., the same image and within category $r_{z}$ from the correlation between the observed and predicted image maps). The results from this simulation indicate that the proposed across-participant pattern similarity method can recover population-level patterns of neural activity attributable individual stimuli.

##### Category Level Similarity

The results from the simulation for the category level similarity measure are shown in Supplemental Figure 3. This figure presents the $r_{z}$ values for each participant (averaged across images) for within and between category component measures derived from correlating the observed beta map with the category maps (i.e., the fixed effect map for the category factor plus the global intercept term). Both the within and between category $r_{z}$ are non-zero, which is expected given they are computed with a shared model term (i.e., the intercept). Critically, the simulated within category $r_{z}$ is higher than the between category $r_{z}$, which indicates that this across-participant approach can also recapture category level neural patterns. Note that both the within and between category measures from the simulated data are higher than that observed data. As discussed with the item similarity simulated measures, this is expected to some extent given that the observed data is likely noisier than the simulated data and the same reasons proposed for the item similarity likely also apply to the category level similarity measures. The critical aspect is that the pattern from the simulated data is like that in the observed data. Thus, overall, the results from this simulation also suggest that the proposed across-participant pattern similarity method can recover population-level patterns of neural activity attributable image categories.

**Supplemental Table 1.** Demographic and neuropsychological test data for young and older adults.

|  | Young Adults | Older Adults | *p*-value |
| --- | --- | --- | --- |
| N | 24 | 24 | - |
| Age | 23.04 (3.46) | 68.92 (3.23) | - |
| Sex | 12/12 | 12/12 | - |
| Education | 15.92 (2.22) | 17.12 (2.23) | .067 |
| MMSE | 29.54 (0.59) | 29.42 (0.93) | .581 |
| CVLT Short Delay – Free | 13.08 (1.79) | 10.83 (2.84) | .002 |
| CVLT Short Delay – Cued | 13.67 (1.81) | 12.33 (2.32) | .032 |
| CVLT Long Delay – Free | 13.54 (2.06) | 10.71 (2.91) | < .001 |
| CVLT Long Delay – Cued | 14.12 (1.62) | 12.33 (2.46) | .005 |
| CVLT Recognition – Hits | 15.42 (0.83) | 15.04 (1.00) | .164 |
| CVLT Recognition – False Alarms | 0.46 (0.66) | 2.67 (2.08) | < .001 |
| Logical Memory I | 30.62 (4.95) | 26.71 (5.09) | .010 |
| Logical Memory II | 28.12 (5.78) | 23.25 (5.72) | .005 |
| Digit Span Total^1^ | 21.04 (4.53) | 17.58 (2.41) | .002 |
| SDMT | 65.38 (13.99) | 47.21 (7.53) | < .001 |
| Trails A (secs) | 21.43 (7.97) | 30.76 (10.77) | .001 |
| Trails B (secs) | 47.54 (19.53) | 69.11 (24.64) | .002 |
| F-A-S Total | 48.29 (10.97) | 45.96 (11.65) | .479 |
| Category Fluency (Animals) | 24.58 (5.67) | 21.08 (4.82) | .026 |
| WTAR (Raw) | 41.42 (3.44) | 43.62 (4.44) | .061 |
| Raven’s (List 1) | 11.08 (.97) | 9.50 (2.23) | .003 |
| Visual Acuity (logMar)^2^ | -.11 (.10) | .06 (.11) | < .001 |

*Note*. Standard deviations are reported in parentheses. The *p*-values were obtained from Welch *t*-tests comparing young and older adults. ^1^Digit span total equals the sum of forward and backward span. ^2^Lower logMAR scores indicate better visual acuity. MMSE = Mini-mental State Exam; CVLT = California Verbal Learning Test II; SDMT = Symbol-Digit Modalities Test; WTAR = Wechsler Test of Adult Reading

**Supplemental Table 2.** Across-participant mean (with standard deviations) of the number of trials by age group, subsequent memory bin, and image category.

|  |  | Age Group | |
| --- | --- | --- | --- |
| Image Category | Subsequent Memory Bin | Young Adults | Older Adults |
| Objects | Source Correct (SC) | 67.5 (30.8) | 66.5 (20.8) |
|  | Source Incorrect (SI) | 28.6 (14.6) | 32.2 (13.7) |
|  | Item Miss | 18.8 (17.5) | 20.2 (13.8) |
| Scenes | Source Correct (SC) | 37.7 (20.1) | 40.5 (17.9) |
|  | Source Incorrect (SI) | 40.2 (12.0) | 43.2 (16.4) |
|  | Item Miss | 38.6 (23.1) | 35.4 (19.6) |

*Note*. Deviations below 120 trials across subsequent memory within each item type and age group result from exclusion of trials for factors such as not responding or responding with the incorrect hand during encoding.

**Supplemental Table 3.** Parameters of the multivariate normal distributions used to create simulated voxel-wise beta maps.

| Parameter | Distribution | µ | σ | Correlation |
| --- | --- | --- | --- | --- |
| Intercept (fixed) | normal | 5 | 1 | n/a |
| Encoding Task (fixed) | multivariate normal | 1 | 1 | .15 |
| Image Category (fixed) | multivariate normal | 1 | 1 | .15 |
| Subsequent Memory (fixed) | multivariate normal | 1 | 1 | .15 |
| Age Group (fixed) | multivariate normal | 1 | 1 | .15 |
| Participant - Young (random) | multivariate normal | 1 | 1 | .15 |
| Participant - Older (random) | multivariate normal | 1 | 1 | .15 |
| Image - Scenes (random) | multivariate normal | 1 | 1 | .15 |
| Image - Object (random) | multivariate normal | 1 | 1 | .15 |
| Random Noise | normal | 1 | 1 | n/a |

*Note.* The random effect beta values were generated separately for young and older participants, and for scene and object images, implying 0 correlation between the two groups. Note that correlations between these groupings were built into the fixed effect beta value.

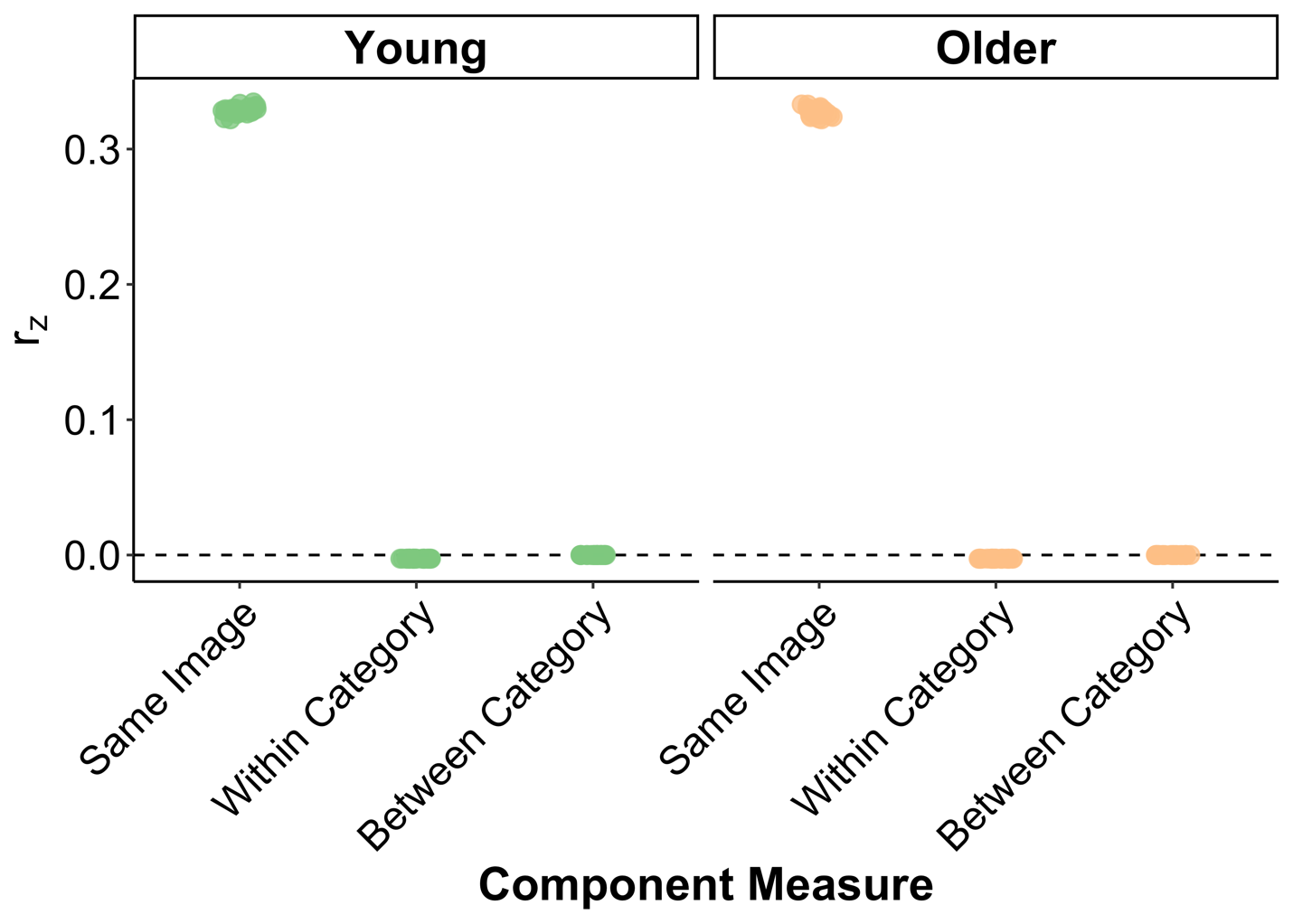

***Supplemental Figure S1.*** The $r_{z}$ values for the same image, within category, and between category components of the item level similarity measure from the simulated data. Note that only the same image and within category $r_{z}$ were used to create the item level similarity measure used in the main text (i.e., same image – within category). The values shown are the average $r_{z}$ across image for each simulated participant to be consistent with how the results are depicted in the main text.

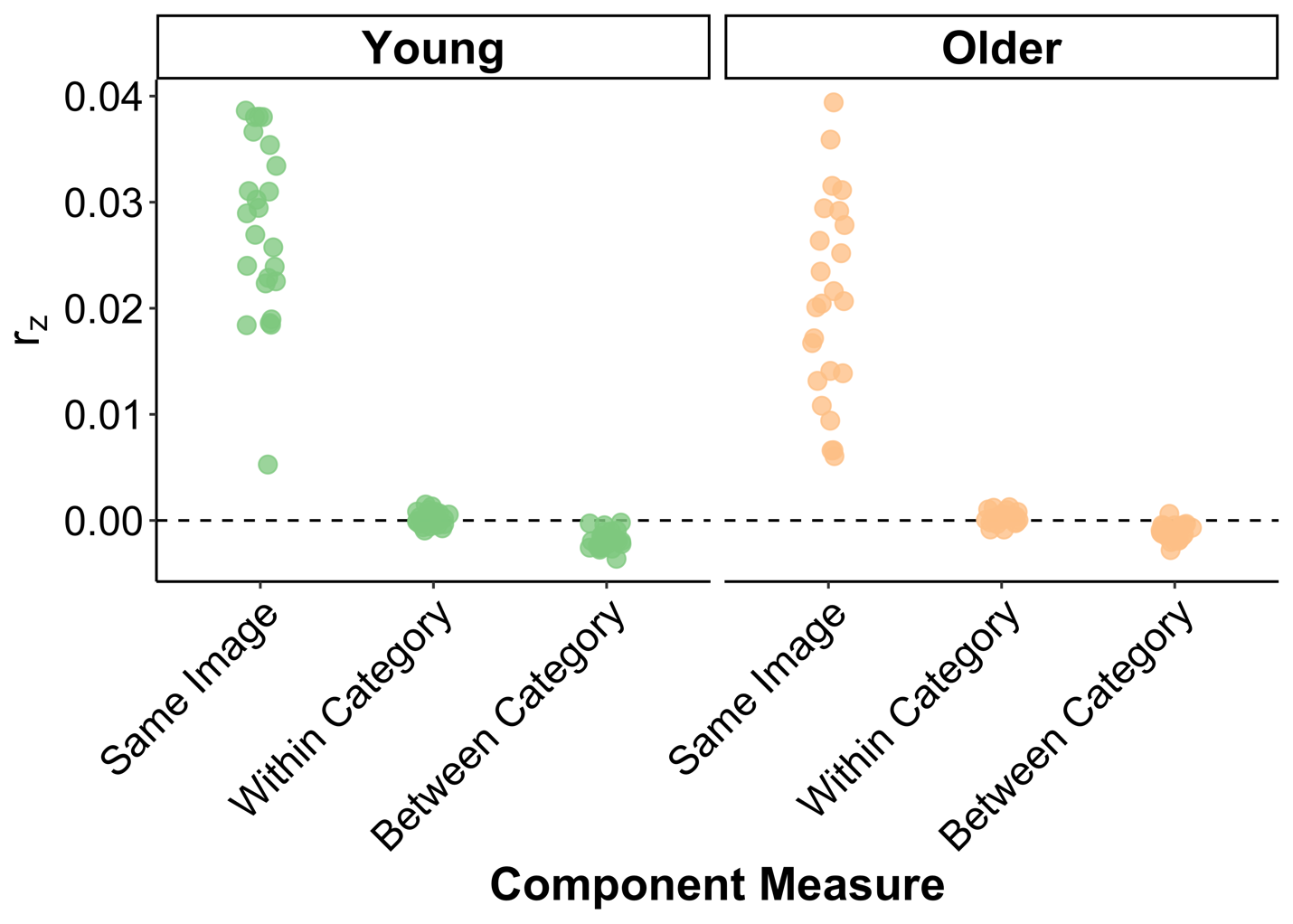

***Supplemental Figure S2.*** The $r_{z}$ values for the same image, within category, and between category components of the item level similarity measure from the observed data. Note that only the same image and within category $r_{z}$ were used to create the item level similarity measure used in the main text (i.e., same image – within category). The values shown are the average $r_{z}$ across image for each simulated participant to be consistent with how the results are depicted in the main text.

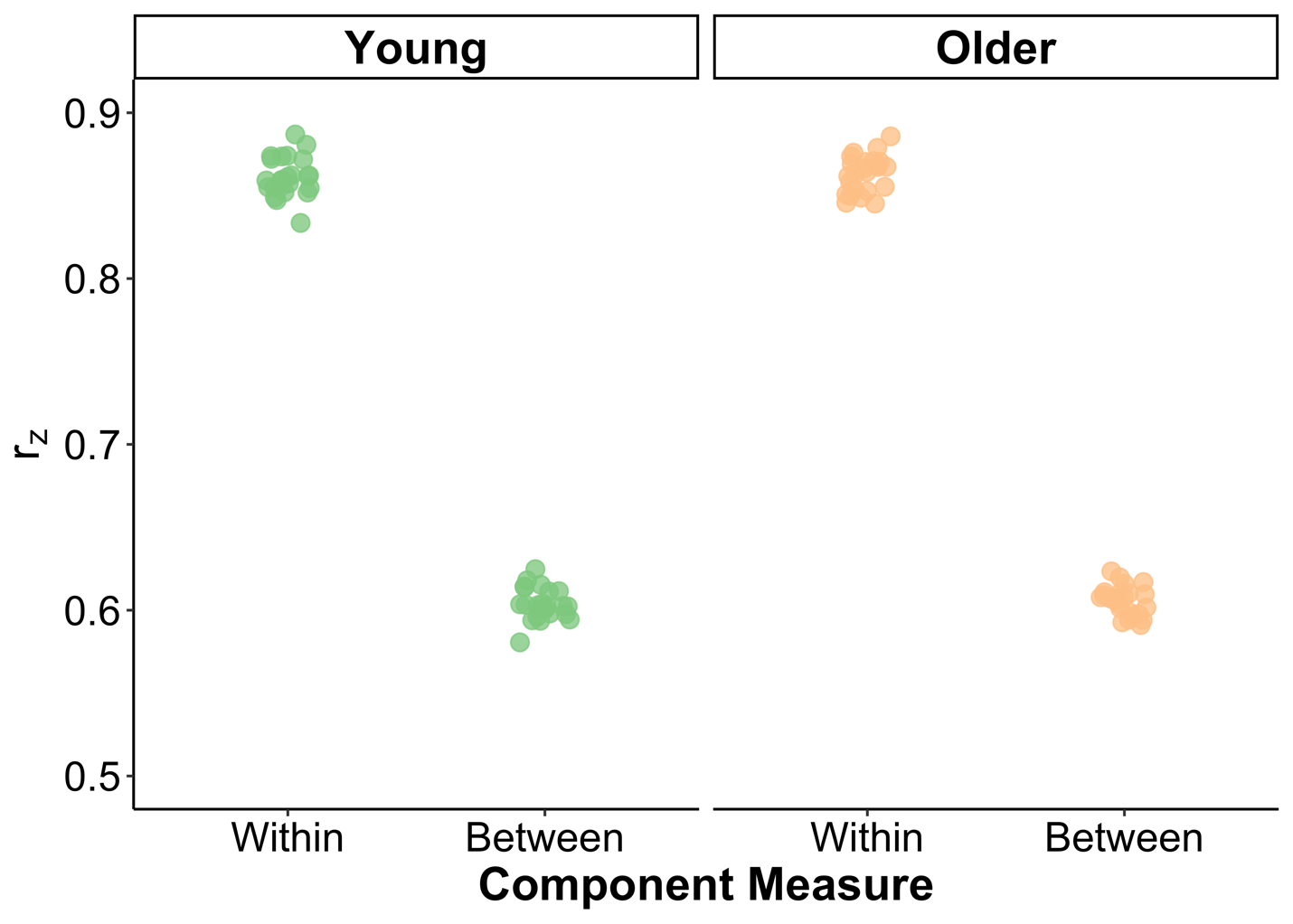

***Supplemental Figure S3.*** The $r_{z}$ values for the within and between category components of the category similarity measure from the simulated data. The values shown are the average $r_{z}$ across image for each simulated participant to be consistent with how the results are depicted in the main text.

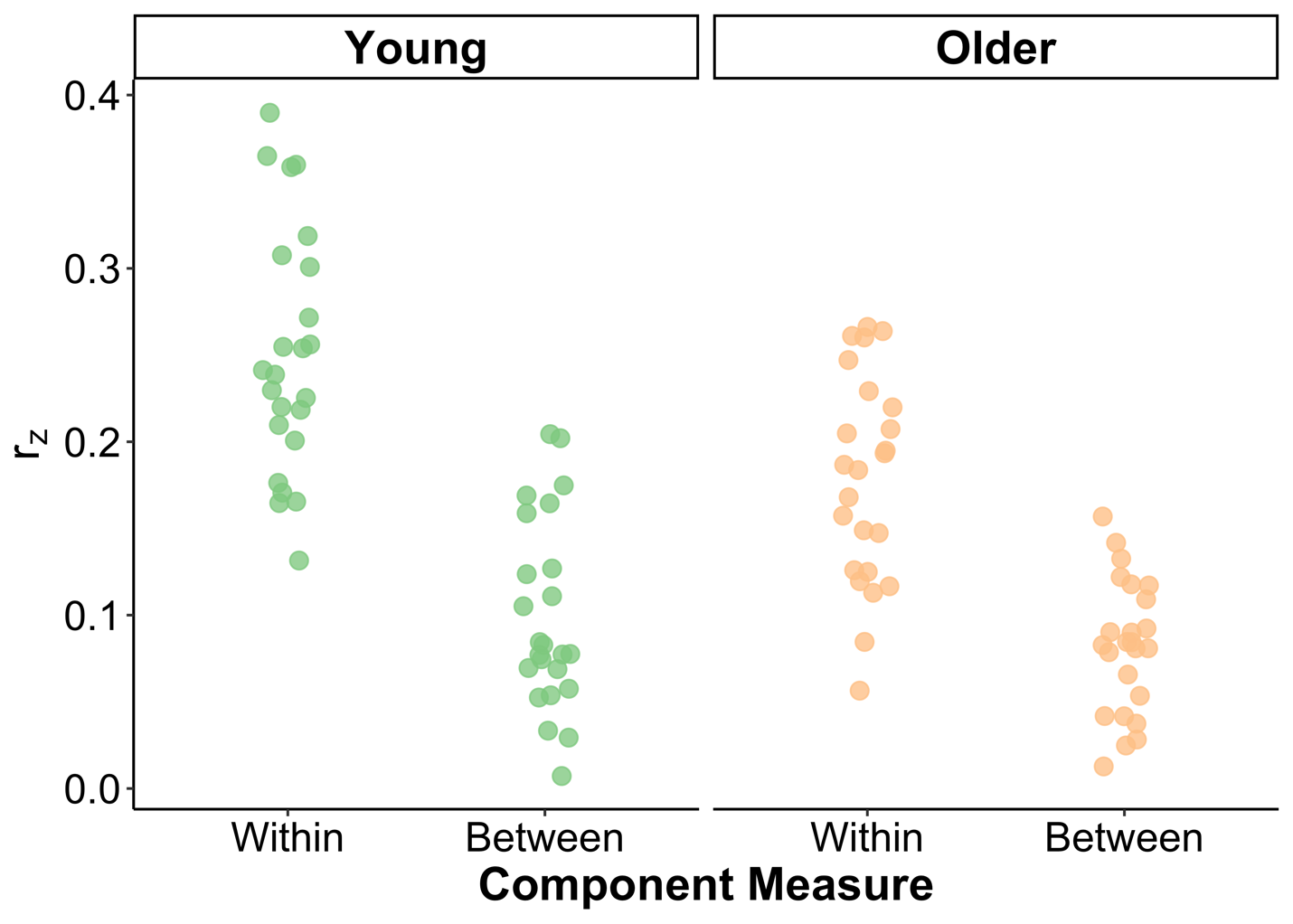

***Supplemental Figure S4.*** The $r_{z}$ values for the within and between category components of the category similarity measure from the observed data. The values shown are the average $r_{z}$ across image for each simulated participant to be consistent with how the results are depicted in the main text.
